# Migratory bird and marine mammal surveillance fails to find evidence for an HPAI H5N1 2.3.4.4b incursion into Australia in 2025

**DOI:** 10.64898/2026.05.07.722556

**Authors:** Michelle Wille, Tobias A. Ross, Robyn Atkinson, David B. Boyle, Maureen Christie, Meagan L. Dewar, Tegan Douglas, Rachael Gray, Birgita Hansen, Rosalind Jessop, Lindall R. Kidd, Ila Marks, Patrick Mileto, Eric Miller, Matthew J. Neave, Sara Ryding, Duncan R. Sutherland, Hui Yu, Marcel Klaassen

## Abstract

The panzootic caused by high pathogenicity avian influenza (HPAI) H5N1 clade 2.3.4.4b has been devastating for animals, globally. Despite global spread, the virus remains absent in Oceania. Herein we report the results of our fourth year of enhanced migratory bird surveillance, coinciding with the spring migration of wild birds in 2025; none of the 847 migratory wild birds or 38 marine mammals were positive for HPAI H5N1, although we did detect LPAI. Surveillance remains a critical tool for HPAI H5N1 response, with early detection and rapid response being critical to mitigate the impacts of this virus on animal, environment and human health.

## Main Text

High pathogenicity avian influenza (HPAI) H5N1 clade 2.3.4.4b has been devastating for animals globally (1, 2). It has caused the destruction of hundreds of millions of poultry(3), an enormous economic impact on dairy farmers in the USA (4), has caused population level declines in marine mammals and avian species (1, 2), and has caused >70 human cases since 2021 (5). To date, HPAI has spread to all continents, with the exception of Oceania (6). Prior to 2024, the most likely route of viral introduction into Australia was considered to be with migratory birds along the East Asian-Australasian Flyway, with millions of shorebirds and seabirds arriving each spring (7). HPAI outbreaks have been ongoing in south-east Asia for decades (3), which is also an important region for incoming migratory shorebirds to stop on their migrations (8). With the long distance spread of HPAI H5N1 through the southern ocean in spring 2024 (9), there is a plausible risk for virus arrival to Oceania with seabirds in the southern Ocean.

Herein we report the results of our fourth year of migratory bird surveillance, coinciding with the spring migration of wild birds in 2025. We investigated 847 migratory birds of the order Charadriiformes and Procellariiformes, and 18 resident Charadriiformes in October– December 2025. Specifically, we captured and sampled short-tailed shearwaters (n□=□254) at a breeding colony on Phillip Island, Victoria, upon their arrival from the northern Pacific. We also sampled 13 Asian-breeding migratory shorebird species at major nonbreeding sites in Exmouth Gulf (n=147) and Peel Estuary (n=83) in Western Australia, Western Port Bay (n=236) and Peterborough (n=17) in Victoria, Canunda (n=19) and Nene Valley (n=□109) in South Australia (Table 1). In addition to migratory and resident birds, we also sampled Australian Sea Lions (*Neophoca cinerea*) in Seal Bay, South Australia (n=38) (Table 1) given the impacts on this animal group in South America and Antarctica. For the sampling animal ethics approvals were obtained from Deakin University animal ethics committee (AEC; B28-2023), Phillip Island Nature Parks AEC (9.2024), Department of Primary Industries and Regional Development, Western Australian Wildlife Animal Ethics Committee (WAEC 25-10-66 and WAEC 22-10-102), University of Sydney Animal Ethics Committee (2025/AE00018) and Federation University Animal Ethics Committee (23-005).

**Table 1:**
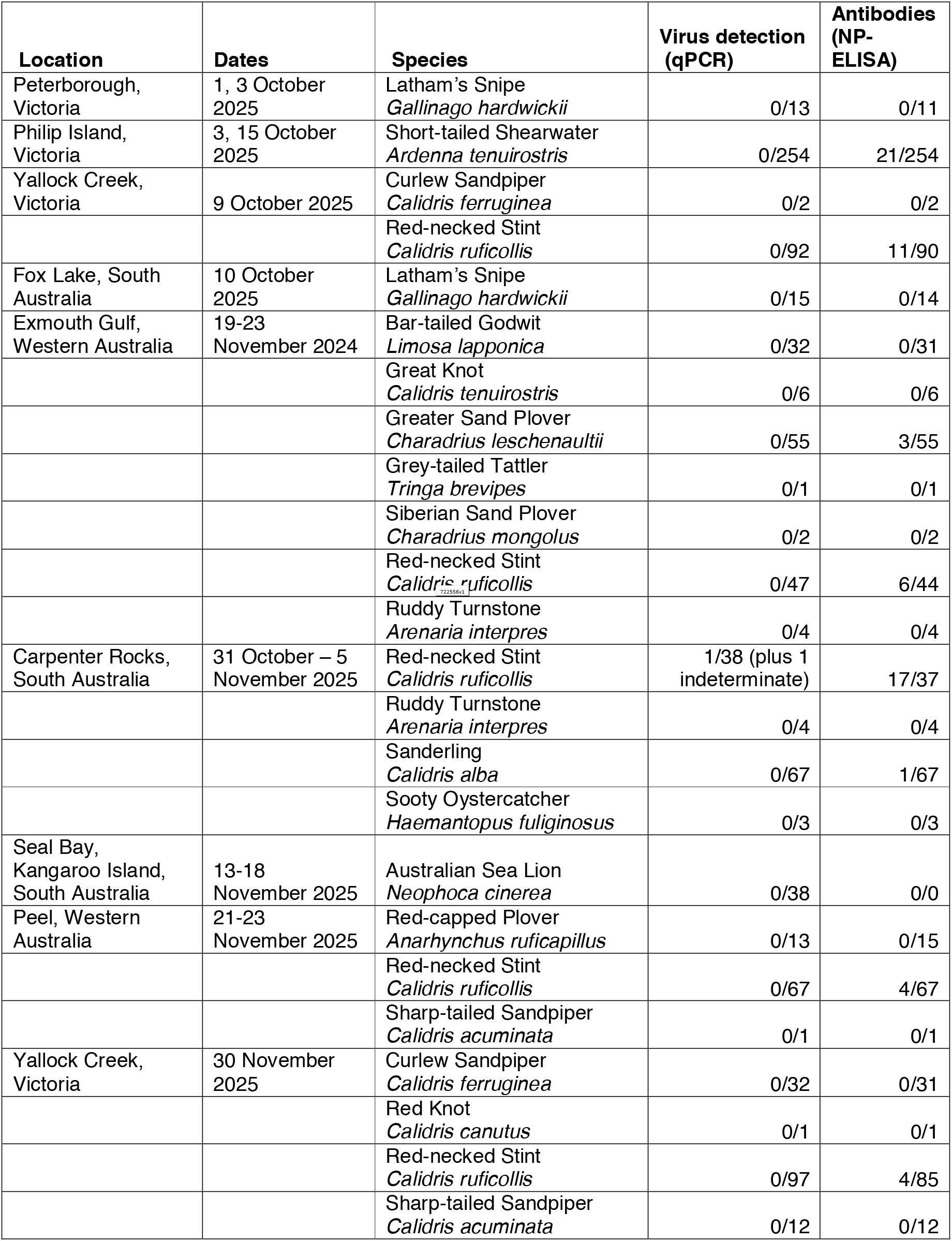
Migratory species targeted for active HPAI surveillance following arrival to Australia after migration from Asia and North America.

Of all 865 avian samples collected, one Red-necked Stint (*Calidris ruficollis*) tested positive (sample ID 21482, Ct=35.8), and one Red-necked Stint was indeterminate (sample ID 21518, Ct=38.7) for influenza A virus by qPCR, following (6). Both samples were negative for H5 and H7 using qPCR (as per 6), and sample ID 21482 was confirmed as H3N7 through full genome sequencing (as per 6). All segments were interrogated using blastn, and phylogenetic trees were constructed using BEAST using approaches and reference sequences reported in (6). The H3N7 virus reported here was similar (HA 99.6%, NA 99.4%) to H3N7 viruses detected in shorebirds in Australia in spring 2024 (6). These viruses are reassortants, with some segments having been in viruses detected in shorebirds >10 years (PA, NP, M, NS), from Australian waterfowl (HA, NA), and from Asian wild birds (PB2, PB1) (Figure 2). All of the samples from Australian Sea Lions were negative for influenza A virus.

**Figure 1.**
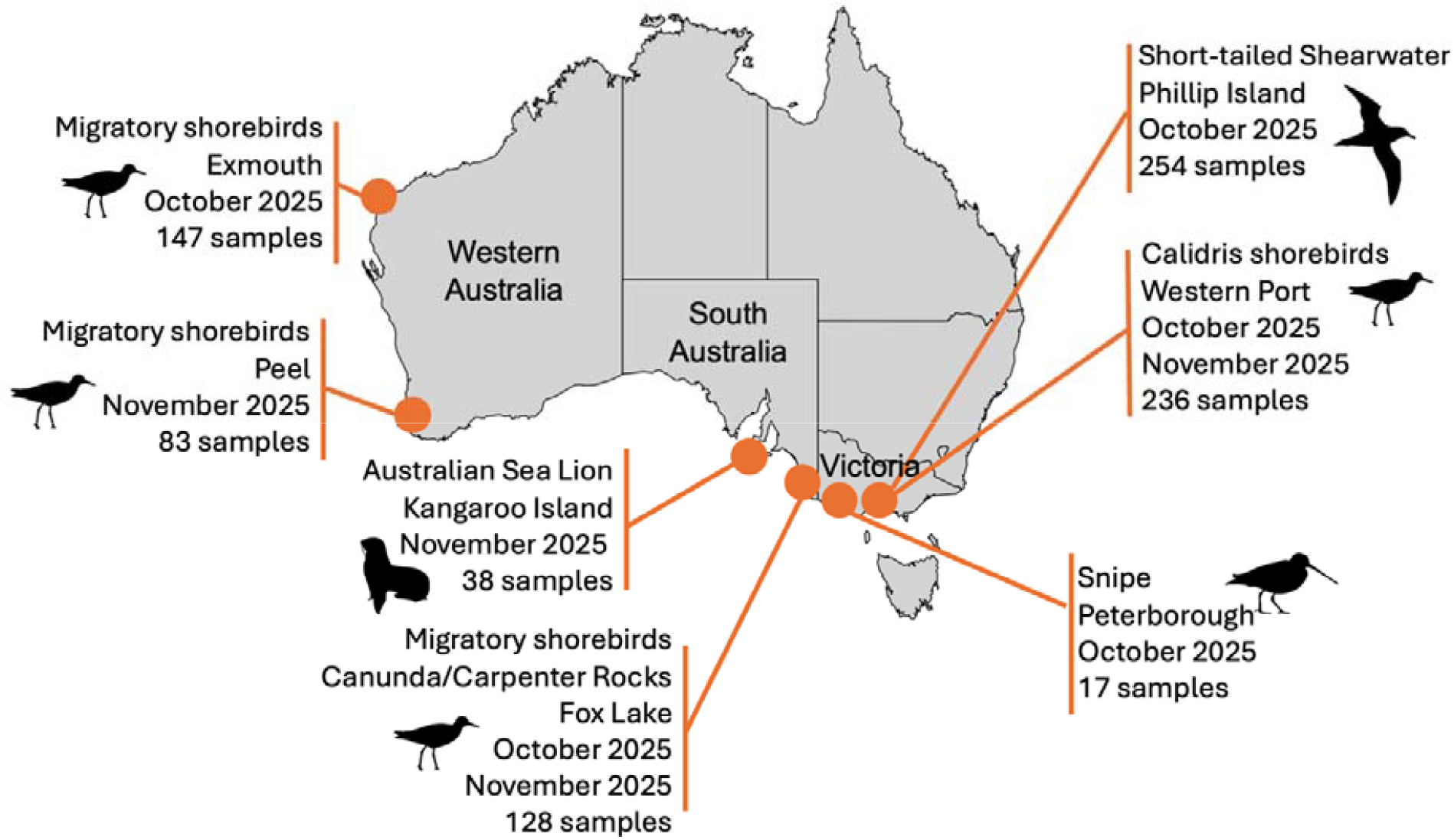
Overview of migratory bird and pinniped sampling between October – November 2025, including locations, main target species, and number of samples collected.

**Figure 2.**
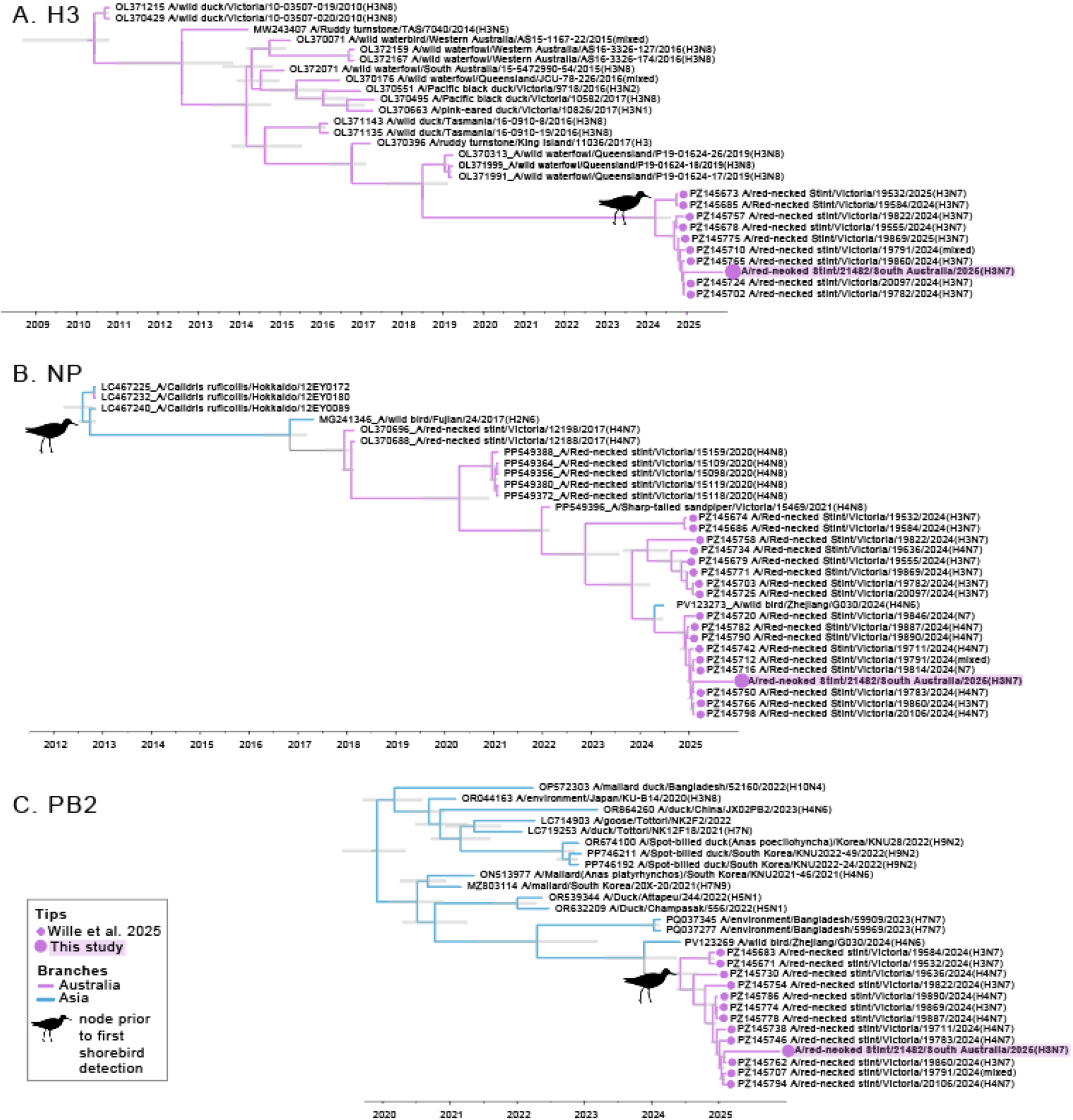
Time structured phylogenetic trees of the (A) HA, (B) NP, (C) PB2 segments of A/red-necked Stint/21482/South Australia/2025(H3N7). All trees include H3N7 and for NP and PB2, also N4N7 viruses from Wille et al. 2025, in addition to top ∼20 blast hits were retrieved from BLAST. Sequences were aligned with MAFFT, clock-like behaviour confirmed using TempEst, and phylogenies were constructed using BEAST 1.10.4, in accordance with Wille *et al*. 2025 (6). Sequences generated in Wille *et al*. 2025 (6) are denoted by a small filled circle, and sequences generated in this study are denoted by a filled circle and shaded box. Node bars are 95% highest posterior density around node height, and scale bar is time in years.

Sixty-seven serum samples tested positive for anti-NP antibodies using a commercial ELISA (given an S/N cut off of 0.6), with Red-necked Stint (42/323) Short-tailed Shearwater (21/254) falling within the previously reported seroprevalence (6) All positive sera samples with enough remaining volume (n=61) were negative on a subsequent hemagglutination inhibition (HI) assay using a lineage 2.3.4.4b candidate vaccine virus A/Astrakhan/3212/2020(H5N8) and a Eurasian lineage LPAI H5N3 now circulating in Australia (A/wild bird/Queensland/P23-02457-0051/2023) (10), following (6). A candidate vaccine virus is a 6:2 recombinant virus on an A/Puerto Rico/8/1934(H1N1)(PR8) backbone with the multi-basic cleavage site removed. In addition to the samples tested here, there is no evidence for HPAI H5N1 in any samples collected for active wild bird surveillance or passive surveillance in Australia, with the exception of Heard Island, an Australian sub-Antarctic Island (11).

Multiple factors determine the likelihood of HPAI H5N1 reaching Australia including the movement patterns of migratory wild birds, the capacity of infected birds to migrate or fly long distances, and the extent of active virus circulation at staging sites used by long-distance migrants. What remains poorly understood is how Australia has continued to avoid an HPAI incursion given continued detections in neighbouring countries and demonstrated ability for animals to move this virus thousands of kilometres.

With the risk of HPAI H5N1, Australia has undertaken considerable preparedness activities including strengthening biosecurity and developing and bolstering response plans (12). Surveillance remains a critical tool for HPAI H5N1 response, with early detection and rapid response being critical to mitigate the impacts of this virus on animal, environment and human health.

## Acknowledgements

We are grateful to all those who assisted with sample collection in the field, specifically S. Salmon (Agriculture Victoria), and all the volunteers of the Victorian Wader Study Group, Friends of Shorebirds SE, Victorian Ornithological Research Group, Phillip Island Nature Parks. We are grateful to P. Eden and S. Ban (Wildlife Health Australia), and A. Breed (Australian Department of Agriculture, Fisheries and Forestry) for ongoing support.

This work was funded by National Avian Influenza Wild Bird Surveillance Program, which receives funding from the Australian Government Department of Agriculture, Fisheries and Forestry and is administered by Wildlife Health Australia. The WHO Collaborating Centre for Reference and Research on Influenza is funded by the Australian Department of Health, Disability and Ageing. ACDP is funded by the Department of Agriculture, Fisheries and Forestry. Exmouth samples were collected with the support of Western Australian Marine Science Institution. Australian sea lion sample collection was supported through Department for Environment and Water, South Australia and Department of Climate Change, Energy, the Environment and Water of Australia. Funding for sampling of Latham Snipe supported by the Renewable Energy Research Initiative under the Department of Climate Change, Energy, the Environment and Water.

## Supplemental Methods

Methods in this study follow Wille *et al*. 2025 (6). We have provided an overview here.

### Sample collection and screening

Migratory shorebirds were captured using cannon-netting and walk-in trapping, and seabirds were captured by picking them up from the ground in the colony. Swabs of pinniped faeces were collected into virus transport media.

From each individual bird, we collected a combined oropharyngeal and cloacal swab into virus transport media. We also collected up to 200µl of blood from shorebirds, and 200µl-400µl from seabirds. Blood was collected from the brachial vein using the Microvette capillary system for serum collection (Sarstedt) and centrifuged ∼10 hours after collection to separate serum from the red blood cells.

### Virus screening

RNA was extracted using the NucleoMag Vet Kit (Scientifix) on the KingFisher Flex platform (Thermo Fisher Scientific). RNA was screened by reverse transcriptase real time PCR (RT-qPCR) for the influenza A virus matrix gene (13) using the SensiFAST Probe Lo-Rox qPCR Kit (Bioline), and a cycle threshold (Ct) cut-off of 40. Positive RNA was referred to the Australian Centre for Disease Preparedness (ACDP) for confirmation, H5/H7 RT-qPCR screening using validated assays, and when possible, full genome sequencing. Full genome sequencing was undertaken on positive samples. We used the dual index library preparation and the Nextera XT DNA Library Preparation kit and 300-cycle MiSeq Reagent v2 kit (Illumina). Sequence reads were trimmed for quality and mapped to respective reference sequence for each influenza A virus gene segment using Geneious Prime software (www.geneious.com) (Biomatters, Auckland, NZ).

### Phylogenetic analysis

Sequences were aligned using MAFFT (14) integrated within Geneious Prime. We utilised the same tree backbones as presented in Wille et al. 2025 as these are the most closely related viral sequences. We confirmed molecular clock-like structure in the data using TempEst (15). Time scaled phylogenetic trees were estimated using BEAST v1.10.4 (16), under the uncorrelated lognormal relaxed clock (17) and bayesian skyline coalescent tree prior (18). We used the SRD06 codon structured nucleotide substitution model (19). One hundred million generations were performed, and convergence was assessed using Tracer v1.8 (http://tree.bio.ed.ac.uk/software/tracer/). Maximum credibility lineage trees were generated using TreeAnnotator following the removal of 10% burnin, and trees were visualised using Fig Tree v1.4 (http://tree.bio.ed.ac.uk/software/figtree/).

### Serology

All sera samples were assayed using the Multi Screen Avian Influenza Virus Antibody Test Kit (IDEXX). An S/N threshold of <0.6 was used. All positive sera samples with enough remaining volume (n=61) were negative on a subsequent hemagglutination inhibition (HI) assay using a lineage 2.3.4.4b candidate vaccine virus A/Astrakhan/3212/2020(H5N8) and a Eurasian lineage LPAI H5N3 now circulating in Australia (A/wild bird/Queensland/P23-02457-0051/2023) (10).

